# The paradox of constant oceanic plastic debris: evidence for evolved microbial biodegradation?

**DOI:** 10.1101/135582

**Authors:** Ricard Solé, Ernest Fontich, Blai Vidiella, Salva Duran-Nebreda, Raúl Montañez, Jordi Piñero, Sergi Valverde

## Abstract

Although the presence of vast amounts of plastic in the open ocean has generated great concern due to its potential ecological consequences, recent studies reveal that its measured abundance is much smaller than expected. Regional and global studies indicate that the difference between expected and actual estimates is enormous, suggesting that a large part of the plastic has been degraded by either physical and biotic processes. A paradoxical observation is the lack of a trend in plastic accumulation found in the North Atlantic Subtropical Gyre, despite the rapid increase in plastic production and disposal. In this paper we show, using mathematical and computer models, that this observation could be explained by the nonlinear coupling between plastic (as a resource) and an evolved set of organisms (the consumers) capable of degrading it. The result is derived using two different resource-consumer mathematical approaches as well as a spatially-dependent plastic-microbial model incorporating a minimal hydrodynamical coupling with a two-dimensional fluid. The potential consequences of the evolution of marine plastic garbage and its removal are outlined.

## I. INTRODUCTION

Every year millions of metric tonnes of plastic are produced on our planet, many of which are eventually dumped into the oceans (Jambeck et al, 2015). Because of the magnitude and damaging consequences of this anthropogenic waste, an increasing concern has been raised along with calls for a transition towards a circular plastic economy (Neufeld et al 2016). Diverse estimates of the amount of plastic have been provided, hovering around 7,000 (Cózar 2014) to 269,000 metric tons (Eriksen 2014) but scarcity of data is still a challenge (Sebille et al 2015) reflecting the difficulties of a precise quantification and as a result many open questions remain (Cressey 2016). Other estimates indicate that even larger amounts of plastic should be expected to be found, forming what some researchers have called the *Plastisphere* (Gregory 2009, Zettler et al 2013). These plastics break down as a consequence of sunlight exposure, oxidation and physical actions of waves, currents and animal grazing (Zettler 2013) producing what is called microplastic, defined as particles that come from 0.33 mm in size to even as small as 20 *μ* m in diameter (Law 2014), is the 90% of the total plastic collected (Eriksen 2014). Predicted estimates indicate that marine plastic waste might increase by an order of magnitude by 2025 (Jambeck et al. 2015).

Most plastic debris can be easily incorporated at all levels within food chains (Davison 2011, Murray 2011, Cole 2013 and Ellison 2007; Farrell 2013, Setala 2014, Galloway et al. 2017). As a consequence, more than 267 different species are affected by plastic debris, including 86% of all sea turtle species, 44% of all seabird species, and 43% of all marine mammal species (Laist 1997). Thereby, the presence of plastic debris in open oceans represents an environmental thread due to its potentially damaging impact on marine ecosystems (Engler 2012) and human consumption of sea food (Rochman et al 2015). It is worth to note that despite all the concerns regarding the potentially damaging effects of plastic, only in the last years it has been possible to obtain reliable estimates, particularly in convergence zones (figure 1a), where most of the debris is found, and the results have been unexpected: the observed amount is much smaller than we would expect, 100-fold of discrepancy (Cózar et al 2014, 2015), providing a strong support to the notion of a substantial loss of plastics from the ocean surface, which it is confirmed by the tremendous loss of microplastics from the global ocean surface between 2007 and 2013 (Eriksen 2014).

**FIG. 1.**
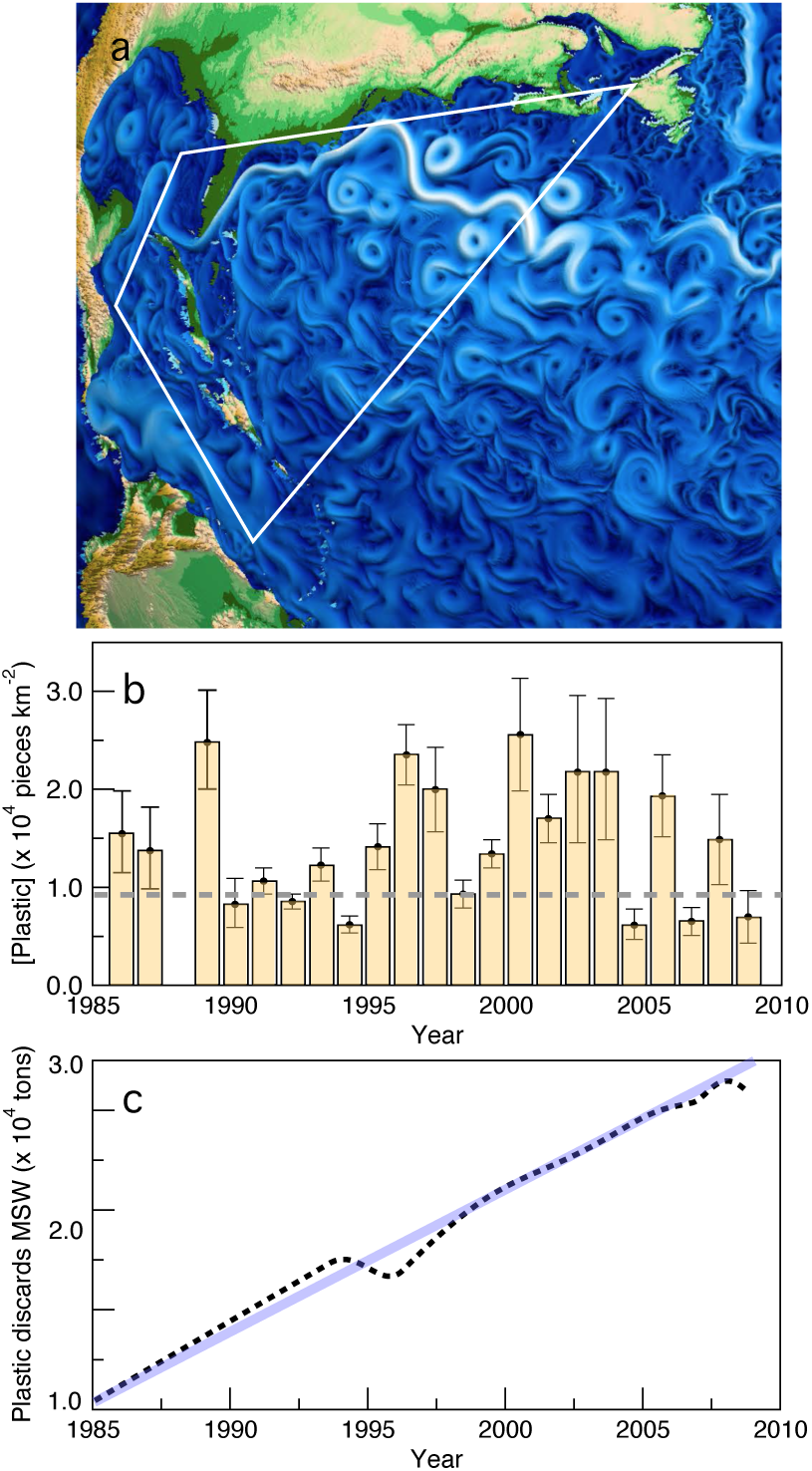
Plastic growing input versus lack of superficial plastic trends. (a) Circulation patterns in the North Atlantic. Here we highlight the area analysed in (Lavender et al, 2010) where marine plastic debris was collected in 6136 surface plankton net tows from 1986 to 2008; the background rendering of ocean currents has been adapted from Wolfram et al 2015. (b) Annually averaged plastic concentration in the region of highest accumulation within the North Atlantic Subtropical Gyre (see Lavender et al 2010) from 1986 to 2008. in (c) we show the concurrent time series of plastic discarded in the U.S. municipal solid waste stream (black points) following an almost linear law.

Remarkably, the analysis of the North Atlantic Subtropical Gyre, a region of highest accumulation of plastics (Law 2010), reveals that the actual amounts of plastic found at the gyre do not exhibit a trend, despite an always increasing rate of plastic entering the ocean, being, in fact, more consistent with a fluctuation around a constant average (Lavender et al, 2010). The analysis of buoyant plastic debris (data available at http://www.geomapa.org) is shown in figure 1b, where the average plastic concentration is shown over a time interval of 22 years. This time series strongly departs from the expected increasing trend given the observed behaviour of plastic discard in the US municipal solid waste stream, also shown in figure 1c. This observation creates a paradoxical situation since it requires some mechanism of plastic removal that cannot be expected by some simple process of linear decay (Wang et al 2016). Even if we take the suggestion of deep sea sinks (Woodall et al 2014) the lack of a growing trend of measured plastic requires an explanation (Franekeer and Law 2015). Why there is no obvious trend present?

In this paper, we propose a potential explanation for this counterintuitive observation based on the assumption that an active and biotic-related process of plastic removal is at work, probably associated to the marine microbial compartment. Whether or not plastics are being degraded by microoganimsm, significantly contributing to the loss of plastic from marine surface, is an open question (Osborn and Stojkovic 2014). In this context of biodegradation, it is known that ocean plastic debris is colonised by a broad range of marine life forms that includes microbes, bryozoans, hydroids or molluscs and other epiphytes (Barnes 2002). Moreover, has been reported that plastic and microplastic can be ingested by different organisms and transported along the trophic chain. Zooplankton can mobilise microplastic (Cole 2013), that is retained in the intestinal tracts for a period of time, in the same way, that occurs in crustaceans (Murray 2011), fishes (Boerger 2010, Davison 2011) or seabirds (Franeker 2011), but eventually, plastic is released again. This is due to the high molecular weight, hydrophobicity and a reduced number of organisms able to metabolise polymers in nature. Nevertheless, is also known that plastics develop biofilms during their residence within the marine environment and that provide habitats for diverse communities of microorganisms, with assemblages of species that differ from those in surrounding seawater (Lobolle 2011, Zettler 2013, Bryant 2016).

Dedicated efforts have allowed the identification of microbial species that seem to be performing active hydrolysis of the hydrocarbon polymers. Furthermore, the possibility that microbes play a key role was confirmed by gene surveys involving small-subunit rRNA, which pointed to hydrocarbon and xenobiotic degrading bacterias (Zettler et al 2013, Bryant 2016). Moreover, a few bacterial strains capable of biodegrading polyvinyl chloride (Shan 2008), polystyrene (Mor and Sivan 2008) or polyethylene (Gilan 2004, Sivan 2006, Balasubramanian 2010, Harshvardhan 2013) have been discovered and more recently, a bacterium that degrades and assimilates plastihas been discovered (Shosuke et al. 2016). Could this evidence be related to the large-scale plastic dynamics?

Our approach is grounded in the formulation of a straightforward hypothesis: the dynamic of plastic debris is affected by a biodegradation process that can be described in its simplest form in terms of a resource-consumer model. We show, both mathematically and numerically, that this pairwise interaction stabilises plastic levels even in the presence of a constantly growing (and thus time-dependent) external input. Given the current lack of parameter estimates, the models presented below are not aimed to fit specific sets of data. Instead, we aim to provide a satisfactory explanation for an unexpected dynamical behaviour and explore its consequences. The model makes specific predictions about the time evolution of consumers and reveals pertinent implications for other problems related to anthropogenic-caused phenomena.

## II. RESOURCE-CONSUMER MODEL

In our computational and mathematical analysis of the problem, we work with two premises concerning the interactions between plastic and the biotic compartment. Firstly, the plastic component will be considered as a homogeneous system with no heterogeneity in terms of composition. Secondly, we do not take into account the multi species nature of the species assemblages interacting with plastic garbage. Instead, we lump together all plastic-degrading species within a single compartment. The basic interactions between plastic (*P*) and a hypothetical consumer (*M*) are summarised in figure 2a. This diagram shows the essential set of assumptions included in our model(s). The simplest version can be formulated by means of a system of two differential equations. The equation for plastic behaviour includes three terms, associated to production, physical decay and active degradation by a microbial consumer, respectively:

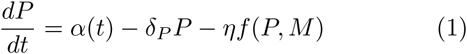

**FIG. 2.**
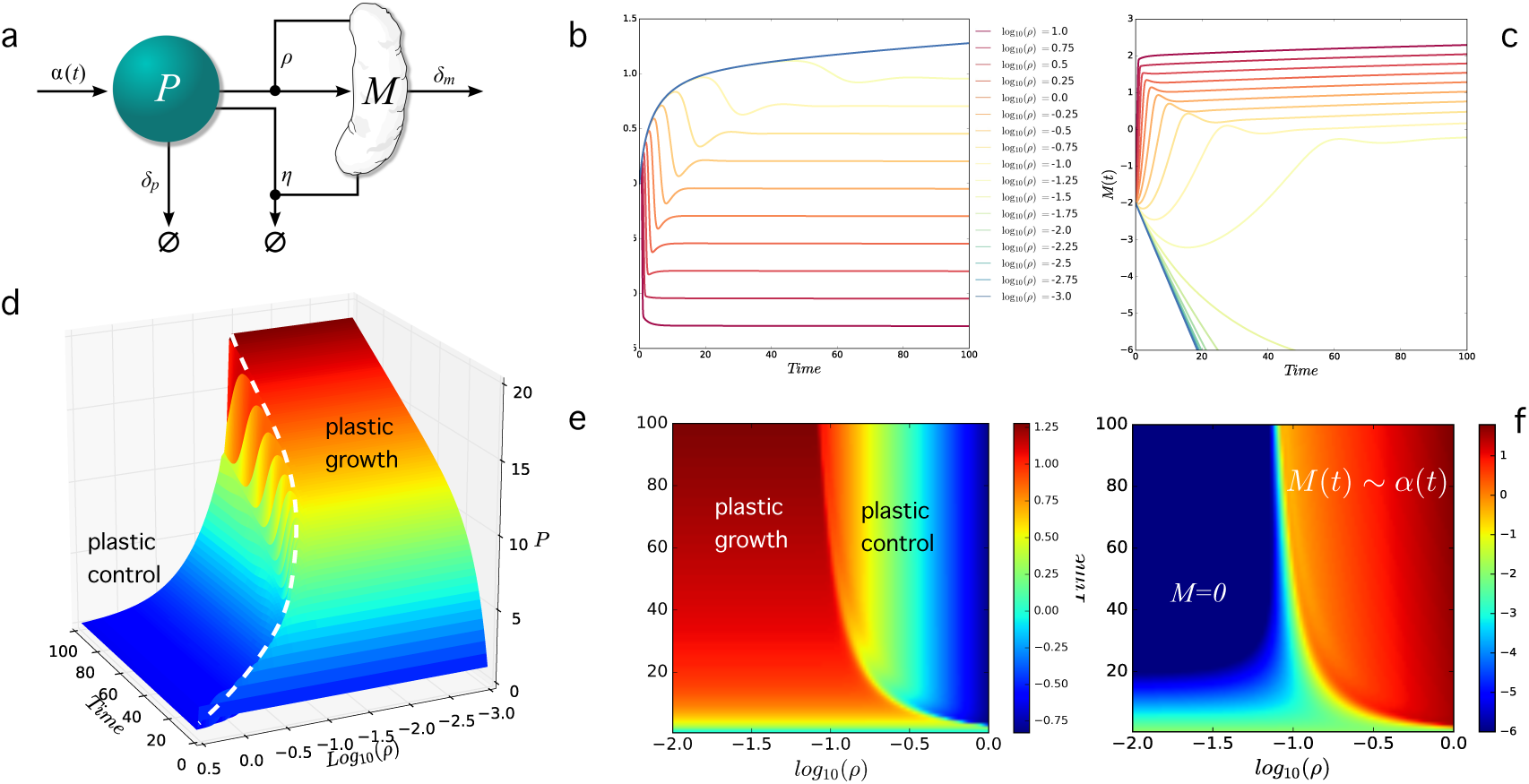
Plastic control by consumer populations. (a) Interactions among plastic (*P*) and microbes (*M*) in the simple resource-consumer model used here. The *M* compartment captures the presence of some hypothetic plastic-degrading species. The time-dependent mathematical model with a linear growth of plastic dumping *α*(*t*) = *βt* shows that, under the presence of the consumer, plastic concentration becomes stabilised to a given fixed value. In (b) and (c) we display the plastic and consumer population time series for our model running over an interval *T* = 500, respectively. The color scale indicates the different values of the parameter *ρ* that measures the efficient of degradation/consumption. The parallel series of consumer levels in (c) follow the input law *α*(*t*) and are observed when the plastic levels achieve constant values. In (d-f) we display these results in a phase space that includes bot the *ρ* parameter as well as the time-dependent dimension in the other axis. Here we use the linear law *α*(*t*) = *α*0 + *βt* with *α*0 = 1 and *β* = 0.01 and an end time *T* = 100. The other parameters arefixed to *δp* = 0.1, *δM* = 0.5 and *η* = 0.20. In (d) the dashed line indicates the points where plastic growth turns into control. Extinction of the *M* population occurs for *M* < 10^*-*6^. In (d-e) we can see that plastic grows when the control by the consumer is strong enough, leading to a constant plastic phase otherwise. The diagram in (f) provides the complementary picture where the consumer gets either extinct or grows following the *α*(*t*) growth function.

Where *α* define the rate of plastic entering in the ocean, *δ*_*P*_ define the rate of physical degradation of plastics and*η* is the efficiency of the consumer degrading plastic. In general, the production will be a time-dependent term and the function *f* (*P, M*) introduces the specific form of the resource-consumer interaction. For the plastic production, we know that it was zero before the late 1040s and starts a continuous trend afterwards, with a constant increase (figure 1c). The second equation includes the dynamics of the consumer, and will read (in its simplest form)

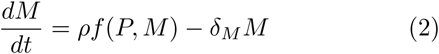

Where *ρ* and *δ*_*M*_ define the growth rate and the death rate of the consumer, respectively. Here the two terms on the right-hand side stand for the growth due to plastic consumption and the death rate of the microbial population, respectively. Let us first consider a linear interaction term, i. e. *f* (*P, M*) = *P M*. In the absence of the consumer (i. e. for *M* = 0) it can be shown (SM) that the plastic concentration follows a time-dependent law:

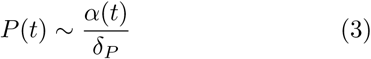

thus growing and accumulating over time. The previous result suggests that we should expect to observe an always increasing amount of plastic resulting from a growing input. If a consumer is present, perhaps we should see a reduced rate of increase, but, as we mentioned before, the paradox emerges from the reported lack of trend exhibited by real data. Here we will show that this is in fact what should be observed if a biotic control from a consumer is at work. Some indication of this possibility comes from the inspection of equation (2) where we can see that the condition *dM/dt* = 0 *allways* gives an equilibrium value *P* ^*\**^ for the plastic (see SM). Below we show how this feature is also maintained when using an extended model incorporating the time-dependent plastic input. These equations are non-autonomous, i. e. they contain explicit time-dependent terms (*α*(*t*)) and thus,the standard stability that assumes constant parameters can no longer be used. Nevertheless, we provide both mathematical and computational proof that a constant (i. e. controlled) plastic density is the expected outcome of the previous model.

Let us first introduce a piecewise approximation by decomposing the input signal *α*(*t*) as a superposition of time intervals Γ_*k*_ (SM) such that, inside each interval, a constant *α*_*k*_ value is used in such a way that our original function is decomposed as

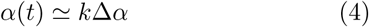

Thereby, for each one of the time intervals, our equations would read:

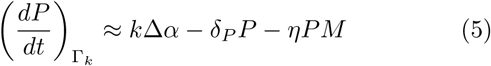

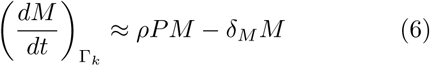

Within each Γ_*k*_ we can use use an approximate stability condition assuming that (*dP/dt*)_*k*_ *≈* 0 and(*dM/dt*)_*k*_ *≈* 0. This leads to a quasi-steady state

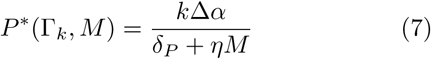

whereas the corresponding steady state for the consumer population is

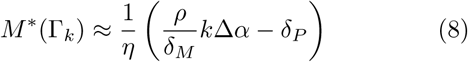

Using these conditions, it can be easily shown (SM) that the dynamics of the consumer populations will grow in time as

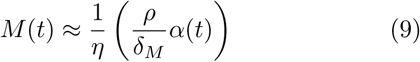

and from this result we can also obtain a dynamical equation for the plastic concentration, namely:

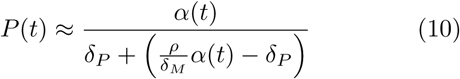

which converges to a constant value of plastic given by

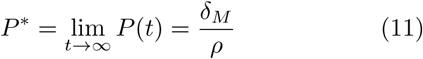

These results not only prove that a saturating (and thus trend-free) plastic debris should be expected. It also predicts a marked increase in the population size of the consumers, which should follow the same trend associated with the rate of plastic dumping over time. Therefore, results are in agreement with the observations of McCormick *et al*, where bacterial assemblages colonising microplastic were less diverse and were significantly different in taxonomic composition compared to those from the water column and suspended organic matter (McCormick 2014). Some numerical examples are shown in figure 2c-f. In Figure 2c we plot the population values *P* (*T*) (black) and *M* (*T*) (red) for a given transient time *T*. The shaded area indicates the domain where no consumer can be sustained and the constant plastic value*P* (*T*) indicates precisely this independence (higher *T*’s would lead to a higher value of these plateaus. As predicted by our model, the microbial compartment rapidly increases while the plastic concentration (now independent of *T* for long times) decays. If the microbe is present, plastic abundance is expected to decay inversely with the efficiency *η* of the consumer (figure 2d) and proportional to its death rate. In other words, the model predicts that the observed plastic abundance is fully determined by kinetic parameters associated with the turnover rates of the consumers. Some examples of the time series associated with the previous plots are also displayed in figure 2e-f.Here small values of *ρ* lead to always increasing plastic levels (and a decay of consumers, black series) where as increasing efficiencies (indicated with different colours) stabilise plastic while *M* grows over time, eventually behaving with the same growth curve for long times (see the parallel lines in figure 2f).

In order to achieve the previous theoretical results, we have introduced a number of simplifications, particularly in relation with assuming a stepwise rate of plastic injection. However, it is possible to rigorously prove that this is a generic property of our resource-consumer model. A general, mathematical derivation based on a detailed analysis of the original non-autonomous system confirms the validity of this result (see SM). Moreover, the results are not changed if other functional responses are introduced. If we consider a chemostat-like model with a Michaelis-Menten saturation term, i. e. *f* (*P, M*) = *P M/*(*K* + *P*), it can be shown that consistently the plastic saturates to a new value *P* ^*\**^ = *δ*_*M*_ *K/*(*ρη* + *δ*_*M*_) whereas anew the microbial compartment scales oncagain, for long times, as *M* (*t*) ∼ *α*(*t*) (see Supplementary Material). Similarly, we should also consider the generalised multispecies model where *S* different types of consumers are introduced, i. e.

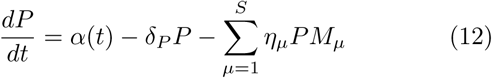

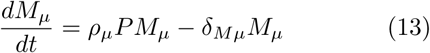

where *μ* = 1*, …, S*. In this case a set of parameters {*η*_*μ*_*, ρ*_*μ*_*, δ*_*Mμ*_} is used. However, it can be shown (see SM) that using the total population *M*(*t*) = ∑ _*μ*_ *η*_*μ*_ *M*_*μ*_ of consumers and the averages 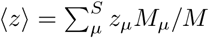 (with *z* = *δ, η, ρ*) we recover the same dynamical equation for the plastic dynamics, namely:

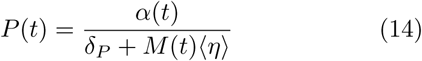

With *M*(*t*) = (〈*δ*_*M*_〉 *α*(*t*))/〈*ρ*〉– *δ*_*p*_)/〈*η* 〉 thus predicting the same outcomes. In our model, as it occurs in resource consumer models with multiple consumers, the competition for the same resource leads to the survival of the best replicator (Smith and Waltman 1995) but the predicted growth law *M*(*t*) ∼ *α*(t) is exactly the same.

## III. SPATIAL DYNAMICS OF PLASTIC-MICROBIAL INTERACTIONS

The consistent results found from these models suggest that our results are robust. However, we have ignored here the potential sources of bias associated to the spatial dynamics resulting from both limited dispersal as well as the physics of plastic movement on the ocean surface. Spatial degrees of freedom can have a major impact on the population dynamics, in particular affecting the stability of the attractors (Bascompte and Solé 1995; Malchov et al 2008). As shown above, our results so far are independent of the underlying details of the interactions among plastic and consumers. Such robustness is further reinforced in this section by considering both spatial degrees of freedom and a toy model including the physics of ocean gyres.

Plastic debris is known to concentrate in gyres (Lavender Law et al 2014, Sebille et al 2015). This process has been approached from several computational perspectives connecting large-scale circulation dynamics with the transport of floating plastic debris (Moore et al 2001; Lebreton et al. 2012, Maximenko et al 2012, Eriksen et al 2014). Here our aim is more limited, since we want to test the potential deviations from the predicted behavior in a hybrid model including currents, discrete plastic items and microbial populations associated to the presence of plastic particles.

The first step in our modelling approach is to generate a vector velocity field **u** on a two-dimensional area (we do not explicitly consider the vertical dimension). To this goal we used a simple hydrodynamic model of fluid dynamics (figure 3a, see SM) while considering two different objects: discrete plastic particles that will behave as rigid bodies and the microbial population, which we consider as continuous. Plastic particles are introduced at a linearly increasing rate, *α*(*t*) defined as the probability per unit time of introducing (random location)a plastic particle. Once a stable vector field has been obtained, it is used as a fixed flow condition (and our results are independent on this particular choice). Plastic particles are introduced in the system at an increasing rate *α*(*t*) at randomly chosen locations (fig 3b). As an initial conditions, we also place a low concentration of microorganisms in each site on the discrete grid.

**FIG. 3.**
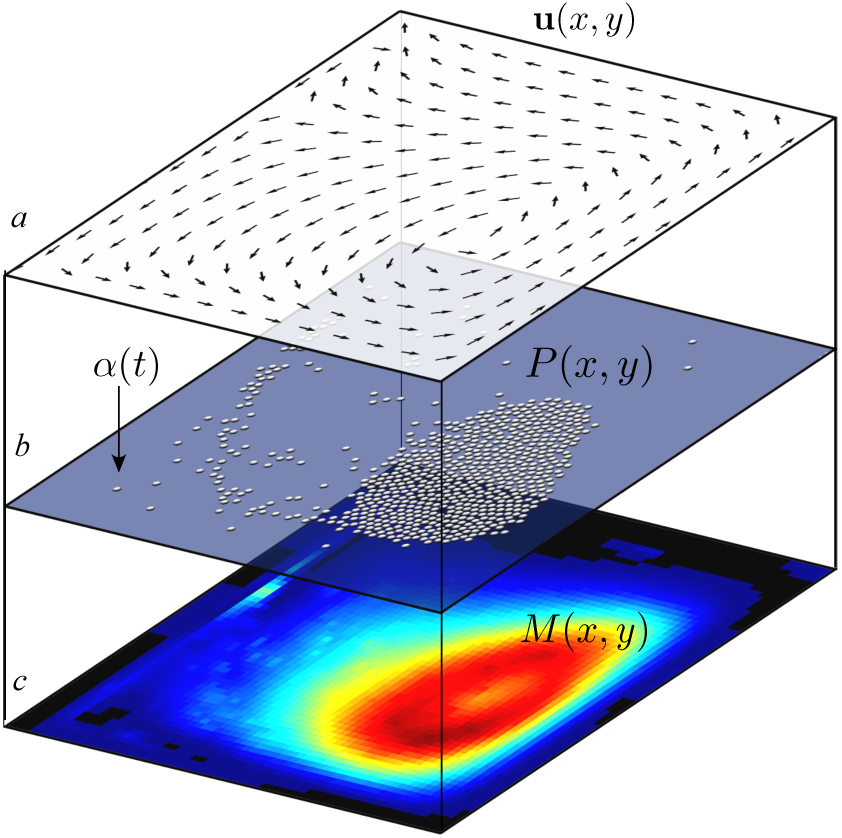
Spatial dynamics of plastic-consumer inter-actions on a gyre. A hybrid model incorporating both the physics of plastic movement on ocean surface and the diffusion-reaction dynamics of microbial consumers is defined by three levels: (a) Surface velocity field generated by a hy-drodynamic model, (b) the set of plastic particles, injected at random locations with some probability *α* and (c) the spatial population of the consumers, defining a continuous field on a discrete lattice.

The corresponding population dynamics term for the microbial compartment (fig. 3c) is then defined, to be affected by both the velocity field as well as the growthdecay dynamics, spatial diffusion and the drift term linked to the velocity field **u**. In terms of a mean field spatial mode, our hybrid model should be able to represent the dynamical process described by the partial differential equation

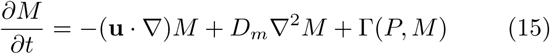

where the three terms of the right hand side incorporate (from left to right) the coupling with the velocity field, passive diffusion (with a diffusion rate *D*_*m*_ and the term Γ(*P, M*), which includes consumer decay as well as plastic-dependent replication (see SM for details). Although plastic particles move on a continuum, the spatial dynamics of *M* (*t*) on our domain *R* will be numerically solved using a discrete mesh Ω = {(*i*, *j*) ϵ *Z*^2^|1 ≤ *i*, *j* ≤ L}. A microbial consumer can be (locally) sustained and experience growth if close to plastic items, while diffusing and being displaced by the plastic particles (acting as a seeding nuclei). Conversely, plastic particles close to consumers will experience a decay in their size, proportionally to the density of consumers in their neighbor-hood.

The hybrid spatial model fully confirms the results predicted by the mathematical model. In figure 4a we show the dependency of the plastic degradation to the efficiency *ρ* of the consumers and how it impacts the time dynamics of plastic, following the same representation as done in fig. 2a. Thereby, while plastic is not controlled for small *ρ* values, it becomes stabilised and remains constant as we move along *ρ* to higher efficiencies. Four spatial snapshots associated to a nearly and late *T* are also shown, both within the controlled (1-2) and uncontrolled(3-4) domains in *ρ* space. In these snapshots we plot both plastic items (gray balls) and the population levels exhibited by the consumer. The contrast between the(2) and (4) snapshots strongly illustrates the two phases, where a clogged system (4) strongly departs from the scattered plastic pieces that remain under the controlled conditions (2) where the consumer population flourishes. In the plastic growth phase, plastic keeps accumulating out of control, whereas in the plastic control phase it becomes constant over time, due to the growth and spread of the consumer through space, which is capable of counterbalancing the increasing input of debris. The connection between the plastic control and the presence of a growing microbial compartment is further illustrated by figures 4b-c. Some examples of the time series of plastic levels displayed by this model are also shown in fig. 4d. Other versions of discrete spatial dynamics with no hydrodynamics, using randomly diffusing particles, give very similar outcomes (results now shown).

**FIG. 4.**
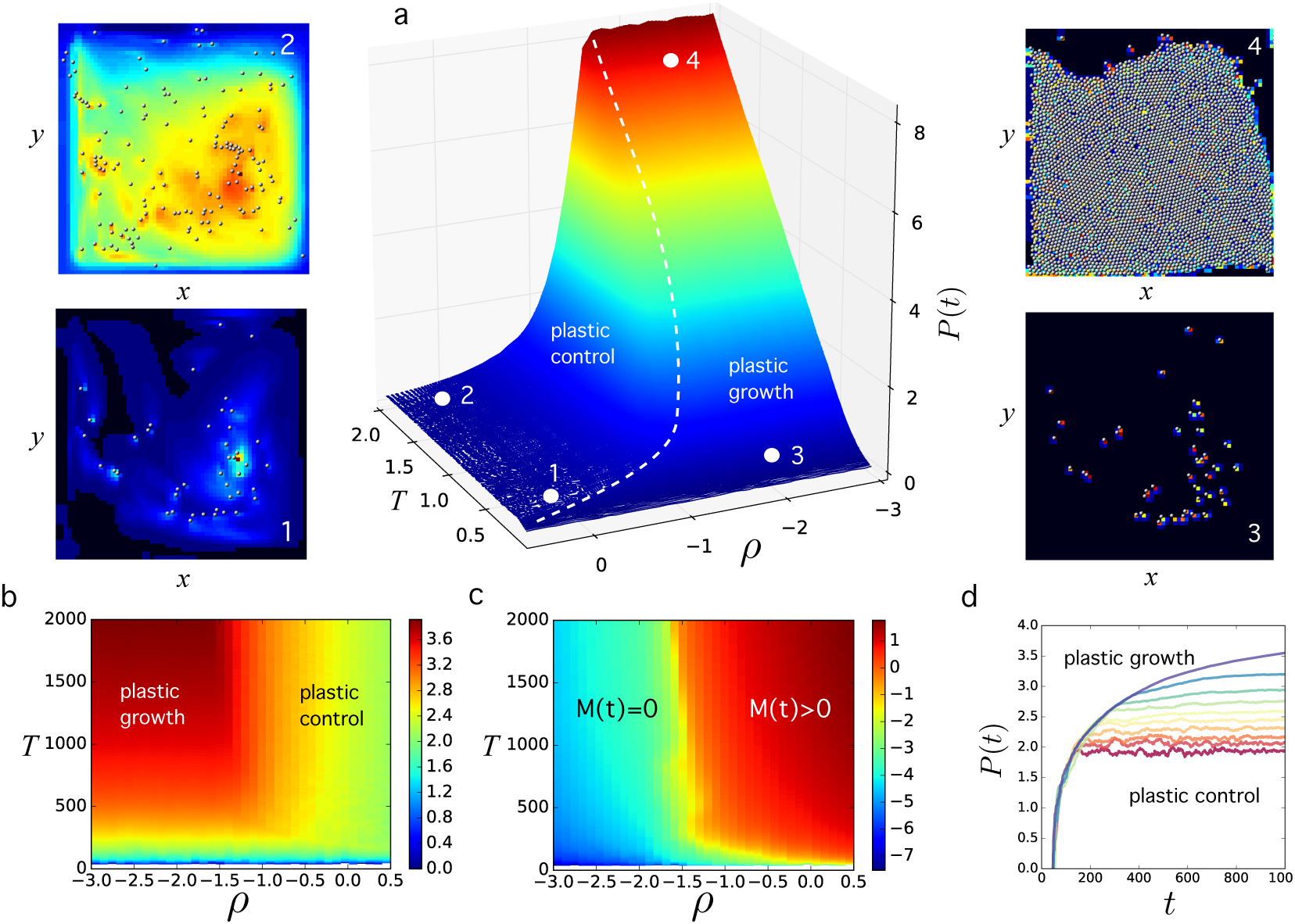
Spatial dynamics of the plastic-consumer model on a gyre. The time evolution of plastic concentration exhibits the same qualitative behavior displayed by the mean field model (fig 2a, logarithmic scale in all axes) with two main phases. The four snapshots (1,2-3,4) indicate the spatial organisation of plastic and consumer concentrations at two given steps *T* for two given *ρ* values. Here plastic (gray balls) is supressed (1-2) and maintained at low concentrations thanks to the growth of the microbial compartment, whereas plastic explodes (3-4) if the efficiency of the consumer is below a threshold value. Two complementary views are given in (b-c) where we can appreciate that plastic control requires the successful growth of consumers. In (d) several time series of the plastic concentration are displayed. The color codes are the same as in fig. 2b-c. The model considers one-vortex (one gyre) system where (a) we display the equivalent diagram of plastic states given in figure 2b-c for the mean field model. The parameters (see SM) are: are *α* = 5 10^*-*3^,*β* = 10^*-*3^, *δp* = 10^*-*2^, *δm* = 10^*-*3^, *η* = 3 10^*-*1^, the initial mass of each plastic particle is set to one.

## IV. DISCUSSION AND CONCLUSIONS

Plastic is a major workhorse of modern economy and a geological indicator of the Anthropocene (Zalasiewicz et al, 2016; Waters et al 2016). Its relevance as a multifunctional material has been increasing since the 1950s along with undesirable ecological impacts. A specially troubling outcome of this is the massive leaking of plastic waste (mostly packaging) to the open oceans. Understanding the dynamics of this enormous Anthropogenic sink is essential to forecast future increases, ecological implications, removal strategies and socio-economic impacts. In this paper, we explored a rather unexpected outcome of the study of ocean plastic debris dynamics, namely: (a) a reduced concentration values (compared to available estimates) and (b) a lack of trend in the total plastic amount despite the known growing trend of plastic damping that should create an easily detectable signal (Lavender et al, 2010; Eriksen et al., 2014; Franeker and Law 2015).

The approach introduced here offers a plausible explanation for both observations. Specifically, we have used a simple resource-consumer model where plastic plays the role of an inert substrate, which is introduced at an increasing rate *α* and degrades linearly (through photochemical and other physical processes) but is also degraded by a consumer. It has been shown that the nonlinearity associated with the interaction between plastic and a biotic partner could account for the observed lack of time-dependent trends in plastic concentration. This interaction can suppress the plastic trend, predicting instead a trend in the consumer population that will match the plastic injection rate. Future sampling efforts should confirm such trend. The robustness of our proposal is illustrated by the consistent results obtained from both the single and multispecies consumer scenarios as well as the hybrid computational approach incorporating the basic spatial interactions taking place in a toy model of ocean gyres.

What are the consequences of these results? Growing evidence supports the presence of microbial candidates for evolved plastic degradation (Shan 2008, Mor and Sivan 2008, Gilan 2004, Sivan 2006, Balasubramanian2010, Harshvardhan 2013, Zettler et al 2013, Shosuke et al. 2016). Our results suggest that hose species associated with plastic debris degradation should exhibit a population boom that correlates with the plastic trend. This could, in fact, help identify new candidates for plastic degradation but also can have consequences for the biodiversity of microorganisms surrounding plastic debris (Chapin III et al., 2000). Constant plastic levels might not be (necessarily) a good news. The rapid degradation of macroplastics predicted by our model also implies a faster transfer from macro-to microplastic that has become a widespread source of potential damage to marine habitats (Laist 1997, Thomson 2004, Browne 2008, Graham 2009, Murray 2011, Farrell 2013, Setala 2014). On the other hand, our model also predicts an expansion of the microbial component, which can include potential pathogenic strains whose dispersal can be favoured by the carrying plastic debris. These hitchhikers can act as plastic degrading agents but also spread across the plastisphere and infect a diverse range of hosts (Kirstein et al 2016; Keswani et al 2016).

What type of strategies can be followed? Some engineering strategies have been proposed to remove macroplastic using gigantic floating barriers (Cressey 2016). The results presented here provide additional insights. The biotic control of plastic could also be achieved using advanced tools from synthetic biology (Khalil and Collins 2010). Specifically, engineered or artificially evolved ecological interactions might be capable of changing the dynamical behaviour of degraded or synthetic ecosystems (Solé 2015, Solé et al 2015). Several design strategies (or *Terraformation motifs*) could combine both genetic and ecological firewalls. One of these designs, to be considered here (Solé et al 2015), would involve a so-called “function and die” design, where the synthetic strain would transiently perform a functional role, such as degrading plastic until niche depletion causes the extinction of the engineered organism. But deep sea microplastic defines an additional niche with a wider and potentially more damaging impact (Woodall et al 2014). Are other species of microbes also exploiting deep sea deposits? If yes, then our results indicate that this might also imply that these wastelands will eventually get reduced to controlled levels, although many known plastics are likely to be difficult to manage by wild type populations. In that case, we could also consider the possibility of designing synthetic strains capable of exploiting (and transforming) deposited microplastics as a main carbon source resource. Finally, another orthogonal possibility should also be considered. Plastic debris is itself a synthetic ecosystem (Solé 2016) capable of maintaining a novel community of organisms. Is the plastisphere worth saving? Could a different engineering strategy capable of stabilising plastic garbage be useful? Both paths need serious consideration as ways of reducing the long-term impact of one of the greatest environmental challenges posed to our biosphere.

## Acknowledgments

We want to thank the members of the Complex Systems Lab for useful comments and discussions. This work was supported by an ERC Advanced Grant Number 294294 from the EU seventh framework program (SYNCOM), by the Botin Foundation-Banco Santander through its Santander Universities Global Division, the Secretaria d’Universitats i Recerca del Departament d’Economia i Coneixement de la Generalitat de Catalunya and by the Santa Fe Institute.

